# Shift in gut microbiota composition and mitigation of diet-induced atherogenesis in mice after prolonged consumption of a traditionally fermented soybean

**DOI:** 10.1101/2025.04.16.649253

**Authors:** Moirangthem Goutam Singh, Himangsu Kousik Bora, Romi Wahengbam

## Abstract

Dietary choices and gut microbiota alterations are linked to the rising prevalence of cardiovascular diseases, including atherosclerosis. Fermented foods, recognized for their health benefits, are known for maintaining cardiovascular health, yet their impact on atherogenesis and associated gut microbiota changes is poorly understood. Here, we showed the restorative potential of long-term dietary supplementation with a traditional Indian fermented soybean, *hawaijar*, in mitigating atherogenic lesions formation and gut microbiota alteration induced by an atherogenic diet in C57BL/6 mice. The diet caused atherogenesis, characterized by increased inflammatory gene expression (IL-1β, ICAM-1, CD4, FoxP3), elevated serum LPS endotoxin levels, and compromised gut health, indicated by increased permeability and decreased expression of tight junction and mucin-producing genes essential for gut barrier integrity (Ocln, Cldn-1, Cldn-4, Muc-2). Metagenomic analysis revealed diet-induced gut dysbiosis, evidenced by an elevated Firmicutes-to-Bacteroidetes ratio, reduced bacterial diversity, and a shift in bacterial composition, including a loss of beneficial *Ligilactobacillus* spp. while pathogenic *Romboutsia* spp., *Escherichia* spp., and *Clostridium* spp. emerged. Conversely, feeding *hawaijar* for sixteen weeks significantly improved atherogenesis, reducing atherosclerotic lesions, serum endotoxins, and inflammatory gene expression. It corrected dysbiosis, restoring gut microbiota to a healthy composition and enhancing gut barrier integrity by reducing permeability and increasing tight junction gene expression. Core beneficial bacteria such as *Phocaeicola sartorii*, *Faecalibaculum rodentium*, and *Akkermansia muciniphila* resurged, alongside the recovery of *Muribaculum gordoncarteri* and *Duncaniella dubosii*, which were lost in the atherogenic condition. Gut eubiosis upon fermented soybean supplementation was also linked with a predicted reduction in major metabolic pathways and a distinct increase in terpenoids and polyketide metabolism. The study highlights the importance of traditional fermented soybean in restoring gut microbiota diversity and gut health against dietary-induced dysbiosis, providing insights into its role in modulating atherogenesis in a gut microbiota-dependent manner.

**Highlights of findings:**

- Metagenomics reveals changes in gut microbiota composition associated with diet-induced atherogenesis and consumption of traditional fermented soybean, *hawaijar,* in C57BL/6 mice.
- Fermented soybean supplementation in atherogenic diet reduces atherogenic lesions, expression of genes associated with inflammatory response (IL-1β, ICAM-1, CD4, and FoxP3), and serum LPS endotoxin level.
- Fermented soybean consumption improves gut barrier integrity, indicated by decreased gut permeability and elevated expressions of tight junction genes (Ocln, Cldn-1, Cldn-4).
- Dietary fermented soybean supplementation promotes eubiosis of atherogenic diet-induced dysbiotic microbiota with a profile characteristic of a normal diet.
- Five core microbiota members, including *Ligilactobacillus* spp., CAG-485 sp002493045 and others, are lost under atherogenic diet regime; however, only *Muribaculum gordoncarteri* and *Duncaniella dubosii* are restored with fermented soybean supplementation.
- Eubiosis of diet-induced dysbiotic microbiota upon fermented soybean supplementation is associated with a predicted reduction in major metabolic pathways and a distinct increase in terpenoids and polyketide metabolism.

**Graphical Abstract:** 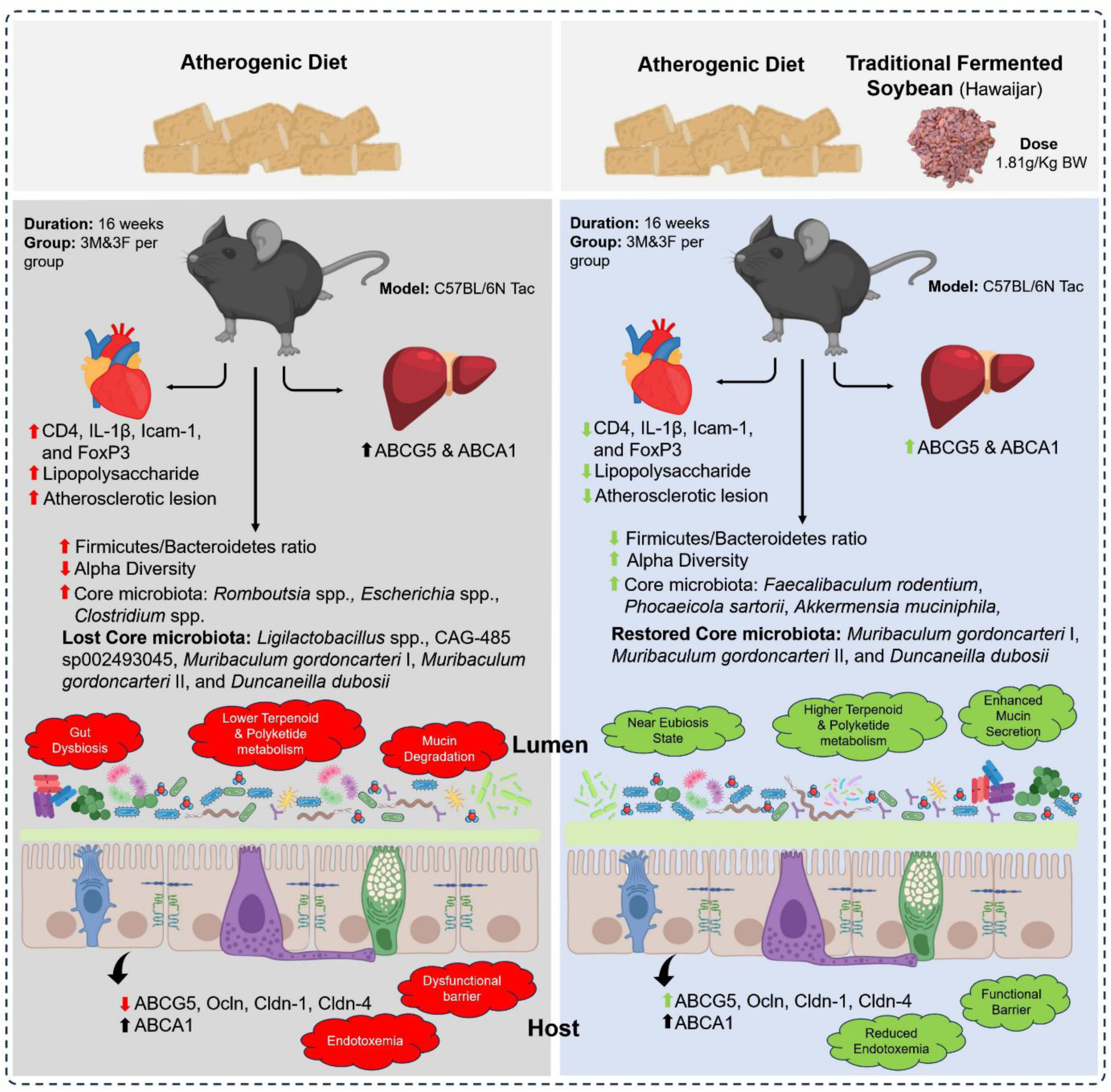

## Introduction

Cardiovascular diseases (CVDs) remain a major public health concern, contributing a considerable share of global death and morbidity. These diverse illnesses cover a wide spectrum of heart and blood vessel disorders, such as coronary artery disease, heart failure, stroke, and hypertension. Atherosclerosis is one of the most important of these. Atherosclerosis, also known as the “silent killer,” quietly infiltrates our arteries, laying the framework for cardiovascular diseases (CVDs). This subtle process involves the gradual buildup of lipid-rich plaques within the artery walls. As these plaques accumulate, they constrict the arterial lumen, hinder blood flow, and pave the way for potentially fatal events including heart attacks and strokes^1^. The hallmarks of atherosclerotic disease are the buildup of cholesterol and the influx of macrophages into the artery wall, which can cause symptoms including myocardial infarction and stroke. The development of atherosclerosis can be influenced by endothelial dysfunction, oxidative stress, hyperglycemia, and chronic inflammation^2^. It has also been demonstrated the dietary lipid phosphatidylcholine is being metabolized into trimethyl amine by the gut microbiota. Trimethylamine has been known to promote atherosclerosis and inflammation in mice. Additionally, choline levels in serum, trimethylamine N-oxide, and betaine serve as biomarkers in predicting CVDs^3^.

Although lifestyle factors and genetic predisposition play significant roles in the development of CVD, new research highlights the potential influence of dietary choices, especially those that contain soy. This high-protein bean contains almost all of the essential amino acids. It is a good source of dietary isoflavones and has low carbs^4^. East Asians consume a lot of traditionally fermented soy foods, which have been linked to beneficial health effects among Asians^5,6^. According to Xu et al.^7^, fermentation increases the digestibility and isoflavone bioavailability of the soybean, improving its functional properties. Other benefits of fermented soybean include immunomodulation, improved nutritional quality and antioxidant activity, and the development of biopeptides that promote health^8^. Fermenting food is a cost-effective and nutrient-dense process with many health advantages^9^. More significantly, food fermentation is typically carried out by bacteria, yeasts, or both, which results in fermented soy meals being an excellent source of pro- and prebiotics ^10,11^. Both conventional and fermented soymilks could have similar effects on increases in *Bifidobacteria* and *Lactobacilli,* with the fermented soymilk having a stronger effect^12,13^. In a population-based cohort study, the dietary intake of soy and *natto*, the well-known fermented soybean of Japan, when taken long-term, was significantly associated with a lower risk of CVD mortality from stroke, underscoring the potential cardioprotective properties of fermented soy foods^14^.

The gut mucosa, the body’s largest immunologically active organ, plays a crucial role in defending the host against invasive pathogens while maintaining a balanced gut microbiota- the diverse community of beneficial and commensal microorganisms that reside in the gastrointestinal tract. These microbes are helpful and perform crucial tasks such vitamin synthesis, xenobiotic chemical metabolism, and nutrition absorption, despite the fact that they influence inflammation and induce infectious disorders^15^. Intestinal bacteria are crucial for controlling the body’s metabolism and inflammation. Given that coronary atherosclerosis is a chronic inflammatory disease and a metabolic disorder, there may also be a close association between gut bacteria imbalance (dysbiosis) and the pathogenesis of atherosclerosis^16^. Furthermore, we hypothesized that the beneficial effect of dietary consumption of fermented soybean in lowering CVD risk ^14^ might be due to the modulation of the gut microbiota, and hence, might ameliorate atherosclerosis.

To our knowledge, no studies have investigated the modulation of dysbiotic gut microbiota in CVD upon consuming traditional fermented foods. The diverse ethnic populations of the North Eastern states of India have traditionally produced and consumed various fermented soybeans as an indispensable part of their diet ^17^. In this context, our study was designed to investigate the impact of prolonged dietary consumption of a traditional Indian fermented soybean, *hawaijar*, in modulating diet-induced gut microbiota dysbiosis and its role in alleviating the atherosclerotic burden. Using the *in vivo* murine model combined with the 16S metagenomic sequencing of fecal samples, we determined the effect of an atherogenic diet and supplementation with the fermented soybean on atherogenic lesions formation, gut microbiota composition, inflammatory response, endotoxemia, and gut barrier health, and predicted the functional pathway or the metabolisms associated with atherosclerosis and gut microbiota changes.

## Materials and Methods

### Experimental Design and Animal Husbandry

All experimental procedures involving animals received formal approval from the Institutional Animal Ethical Committee (Approval No. NEIST/IAEC/2021(1)/001). Stringent ethical guidelines were followed throughout the study. We procured 8-10-week-old C57BL/6N Tac mice, weighing 20-25 grams, from Vivo Bio Tech Ltd., Telangana, India. The mice were housed in controlled environmental conditions, temperature at 25±2°C and humidity 70-80% with 12h light/dark Cycle. Prior to any experimental interventions, a three-week acclimatization period allowed the animals to adapt to their new environment. During this period, *ad libitum* access to both feed and water was ensured. Following acclimatization, we randomly assigned the mice to four experimental groups, each comprising 3 males and 3 females (*n*=6). The normal diet group (Nc) received a standard chow diet, Altromin 1320 (Altromin, Germany). The atherogenic diet group (At) was fed a modified atherogenic diet, Altromin C 1061, consisting of 20% fat, 1.25% cholesterol, and 0.5% Na-cholate. The NcFSB received standard chow diet with the supplementation of the traditional fermented soybean (FSB) from Manipur, known as “Hawaijar” (Figure S5). Finally, the group AtFSB received the atherogenic diet along with FSB supplementation. The administered dose of FSB was precisely 1.8107 g/kg body human equivalent weight (Figure S1a). The animal experiments spanned 16 weeks, during which we closely monitored health, activity, and any potential effects arising from the dietary interventions. After the study period, the mice were euthanized, and various tissue samples were collected and preserved either in formaldehyde or RNAlater for downstream assays.

### Body weight and feed consumption

Each experimental cage received an initial allocation of 20 grams of feed. On the subsequent day, we quantified the remaining feed by precisely measuring the weight of unconsumed portions. Weekly measurements of individual animal weights were conducted. These assessments provided critical insights into growth trajectories and potential variations. The health status of each animal was vigilantly monitored throughout the study period. Any signs of distress, illness, or abnormal behaviors were promptly noted.

### Microbiota composition in fecal samples

Weekly collected fresh feces samples were snap-frozen and immediately preserved at -80°C until processed. Metagenomic DNA was isolated with the QIAampPowerFecal Pro DNA Kit (QIAGEN, Germany), as per the manufacturer’s instructions. Samples were eluted to a volume of 100µl with the elution buffer (1X Tris.Cl, pH 7.4) and stored at -80°C. 16S-V4 region was amplified in a 25 μL reaction volume using 0.75X PlatinumII Hotstart 2X Master Mix (Invitrogen), 0.2μM primers (forward: TCGTCGGCAGCGTCAGATGTGTATAAGAGACAGGTGYCAGCMGCCGCGGTAA, reverse: GTCTCGTGGGCTCGGAGATGTGTATAAGAGACAGGGACTACNVGGGTWTCTAAT) (Macrogen, South Korea), and 5μL template DNA, with the following PCR conditions: initial denaturation at 94°C, 3mins; cycling denaturation at 94°C, 45s; annealing at 50°C, 60s; extension at 72°C, 90s for 25 cycles; final extension at 72°C, 10mins; 4°C, hold. The reactions were set up in duplicates to reduce the technical bias. The replicate PCR products were pooled and cleaned up using AMPure XP beads (Beckman Coulter, USA). The purified 16S-V4 PCR products were indexed with Nextera XT v2 index kit, as per manufacturer’s instruction (Illumina,USA) and the indexed individual samples were normalized and pooled to give a 4nM library. The 4nM library was denatured, diluted to 7 pM and spiked with 25% of PhiX to maintain a balanced nucleotide diversity. The sequencing was run using 600 μl of 7 pM spiked-in library in an Illumina MiSeq platform using MiSeq Reagent Kit v3 and 2x250 bp paired-end reads. The demultiplexed fastq files were analyzed using the QIIME2 (v2023.5)^18^. DADA2 plug-in was used to filter and trim reads based on the quality score, merge pair-end reads, filter chimera, identify amplicon sequence variants (ASV), and quantify their abundance^19^. Taxonomy assignment was done against Greengenes 2.0 database^20^. After the DADA2 denoising (Table S1), the non-chimeric features were OTU-clustered at 99% similarity based on the Greengene2 database. The OTUs were taxonomically classified using Naïve Bayes_trained_unweighted 16S-V4 Greengene2. Mitochondria and chloroplast DNA, singleton, doubleton, less than 0.01% of total frequency and less than 1% abundance across the samples were filtered during the analysis (Table S2). The feature ASV table, taxonomy file and metadata file were utilized to further analyze and generate different visualizations in R, MicrobiomeAnalyst2^21^ and OriginPro.

### Atherosclerotic lesion assessment

A 0.5% Oil Red O stock solution (HiMedia, India) was prepared in 100% isopropanol. The stock solution was diluted 1:1 ratio with ddH_2_O to make working solution. The formaldehyde-fixed aortic arch was washed twice with 1X PBS (pH 6.5) in 1.5 ml tubes for 2 minutes with a gentle vortex followed by 70% ethanol in 1.5 ml tubes. 1ml of freshly prepared Oil Red O stain was added and incubated in a thermo-mixer at 24 °C for 1 hr. The stained aortic arches were transferred to a new 1.5ml tube and washed with 70% ethanol, followed by 1X PBS. The stained tissues were observed in a stereo stereo zoom microscope, and images taken. The observed images were analyzed for the percentage of the atherosclerotic lesions using ImageJ software.

### Biochemical analysis of serum samples

Blood samples were collected via retro-orbital plexus under isoflurane anesthesia. Blood samples were allowed to clot and centrifuged, and the obtained serums were preserved at −80°C until analysis. Total cholesterol and low-density lipoprotein cholesterol (LDL-C) levels were determined using commercially available kits from Peerless Biotech, India. Plasma lipopolysaccharide (LPS) levels were quantified using the ToxinSensor^TM^ Chromogenic LAL Endotoxin Assay Kit (GenScript, USA). Relative intestinal permeability was measured using the non-metabolizable macromolecule Fluorescein Isothiocyanate (FITC)-Dextran (4kDA, Sigma-Aldrich, USA) following the protocol described by Thevaranjan et al.^22^

### Gene expression profiling

Total RNA was extracted using the Monrach® Total RNA Miniprep kit (New England Biolabs, USA) from the whole aorta, liver, and proximal colon preserved in RNAlater (Sigma-Aldrich, USA). The cDNA (20 μl) was synthesized from 100ng RNA using LunaScript® Super Mix Kit (New England Biolab) and diluted 2X with nuclease-free water to set up the qRT-PCR reactions in a 10μl reaction volume using 1X PowerUp^TM^ SYBR^TM^ Green Supermix (ThermoFisher Scientific, USA) and 400nM gene-specific primers (Table S3). The reactions were run on a QuantStudio 5 Dx Real-Time PCR System (Thermo Fisher Scientific). The relative gene expression was calculated by the ΔΔCt method using β-actin as the internal control. The fold change was calculated as 2^-ΔΔCt^.

### Functional prediction

Functional prediction of gut microbiota was carried out using PICRUSt2^23^ based on the 16S-V4 sequence data. Fasta file of representative sequence (.fna) and ASV feature table (.biom) were used to generate the KO abundance file. Differential abundance analysis was performed using the “LinDA” method. The top 50 significant pathways were annotated using the KEGG database. The major functional categories were assigned based on the BRITE hierarchy in the KEGG database. Abundance mean and standard error were calculated and log transformed.

### Statistics

All statistical analyses were performed in OriginPro 2024b (learner edition) and R(4.4.0). All the tests statistics were included in the figure legends. The mean represents the central point of the error bars in the figures, and the standard error of the mean is represented by the top and bottom edges. Significant value was calculated using either an unpaired *t*-test, a two-way ANOVA, or a Kruskal-Wallis test, with *p* values adjusted using the Benjamini-Hochberg method. Non-significant *p* > 0.05; \**p* ≤ 0.05; \*\**p* ≤ 0.01; \*\*\**p* < 0.01.

## Results

### Fermented soybean supplementation reduces atherogenic diet-induced atherosclerotic lesions

Atherosclerotic lesions were quantified using ImageJ software (Figure 1a), with the lesion area presented as a percentage (Figure 1b). The highest percentage of atherosclerotic lesions was observed in the atherogenic diet-fed (At) group, followed by the fermented soybean *hawaijar* (FSB)-supplemented atherogenic diet-fed (AtFSB) group, while the normal chow diet-fed (Nc) group had a lower lesion area, and the FSB-supplemented normal diet-fed (NcFSB) group exhibited the smallest lesion area. Marker genes of vascular inflammation, including CD4, IL-1β, ICAM-1, and FoxP3, were significantly upregulated in the At group (Figure 1c). Additionally, serum total cholesterol and LDL levels were elevated in the At group than the Nc group (Figures 1d and 1e), potentially contributing to the heightened expression of inflammatory genes. However, FSB supplementation led to a reduction in both the atherosclerotic lesion area and the expression of the inflammatory genes. No significant changes in serum cholesterol or LDL levels were observed in the AtFSB and NcFSB groups.

**Figure 1.**
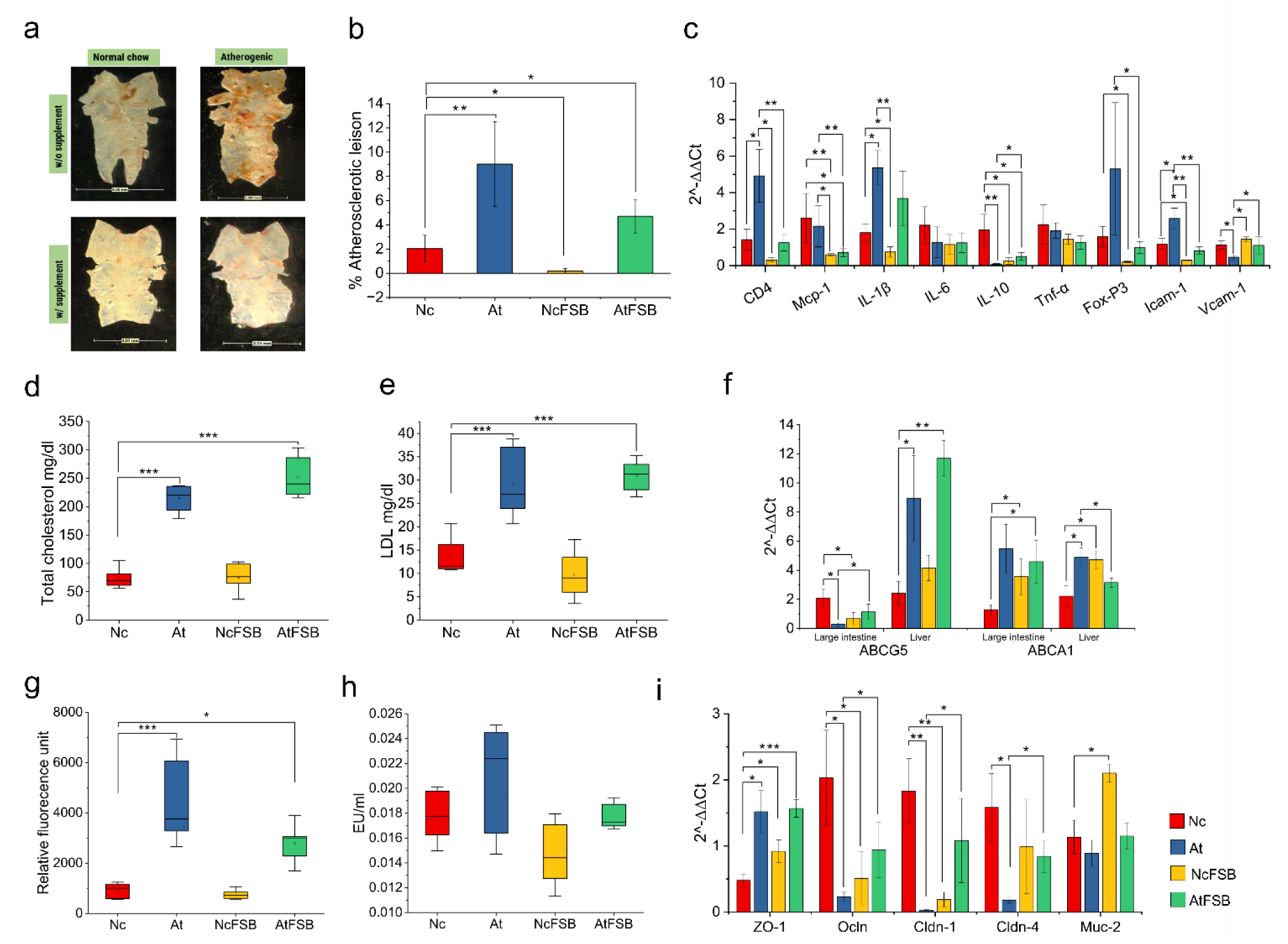
Effect of atherogenic diet in the formation of atherogenic lesions, inflammatory genes expression, total cholesterol and LDL levels, and gut barrier integrity, and comparison with FSB supplementation. (a) Image showing the Oil Red O stain of the atherosclerotic lesion in the arch of the aorta. (b) Bar chart of % atherosclerotic lesion of the experimental groups. (c) Bar chart of the fold change of genes associated with inflammatory response during atherogenesis. (d) (e) Boxplots showing the levels of total cholesterol and LDL in the serum, respectively. (f) Bar chart shows the fold change expression of cholesterol transporter genes, ABCG5 and ABCA1, in the liver and large intestine of each group. (g) Boxplot showing the level of serum FITC-Dextran (4kDa) in each group. The units are expressed in relative fluorescence units. (h) Boxplot of the level of serum LPS endotoxin in each group. (i) Bar chart showing the fold change expression of tight junction genes in the large intestine. Significance was calculated using Kruskal-Wallis. Non-significant p > 0.05; *p ≤ 0.05; **p ≤ 0.01; ***p < 0.01.

Cholesterol transport genes, ABCG5 and ABCA1, were more highly expressed in the liver of the At group than the Nc group (Figure 1f). Conversely, ABCG5 expression in the large intestine was reduced, while ABCA1 expression remained elevated in the At group. FSB supplementation influenced gene expression, increasing ABCG5 levels in both the liver and large intestine of the AtFSB group. Although ABCA1 expression decreased in the AtFSB group, this reduction was not statistically significant. In the NcFSB group, both ABCG5 and ABCA1 expression were upregulated in the liver and large intestine.

### Supplementation of fermented soybean improves gut barrier health

The gut permeability assay indicated that the level of the non-metabolizable macromolecule FITC-Dextran in the At group were significantly higher than the Nc group, suggesting increased intestinal permeability in the At group (Figure 1g). This observation was corroborated by the elevated serum LPS endotoxin levels in the At group (Figure 1h). Although endotoxin levels were higher in At than in Nc, the difference was not statistically significant due to inter-individual variability. Increased intestinal permeability in the At group was further supported by the downregulation of key tight junction genes (Ocln, Cldn-1, and Cldn-4) and mucing-producing gene (Muc-2) (Figure 1i). The supplementation of atherogenic diet with FSB (AtFSB group) reduced intestinal permeability and serum endotoxin levels. This stabilization effect was also reflected in the AtFSB group where the expressions of tight junction and mucing-producing genes were upregulated as compared to the At group.

### Atherogenic diet-induced gut microbiota diversity loss and shifts in its composition

We explored the changes in gut microbiota composition induced by the atherogenic diet. In the Nc group, Bacteroidetes dominated the microbial community during weeks 1-4, followed by Firmicutes, Cyanobacteria, Actinobacteria, Desulfobacterota, Deferribacterota, Proteobacteria, and Verrucomicrobiota. However, during weeks 5-8, a notable shift occurred, with increased relative abundances of Firmicutes, Proteobacteria, Desulfobacterota, Deferribacterota, Cyanobacteria, and Actinobacteriota, while Bacteroidetes and Verrucomicrobiota decreased. By weeks 9-12, Bacteroidetes, Actinobacteriota, Desulfobacterota, and Proteobacteria showed an increase, whereas Firmicutes, Deferribacterota, and Verrucomicrobiota declined. In the final phase, from weeks 13-16, Firmicutes maintained their dominance, followed by Bacteroidetes, Desulfobacterota, Actinobacteriota, Cyanobacteria, Deferribacterota, Proteobacteria, and Verrucomicrobiota (Figure 2a).

**Figure 2.**
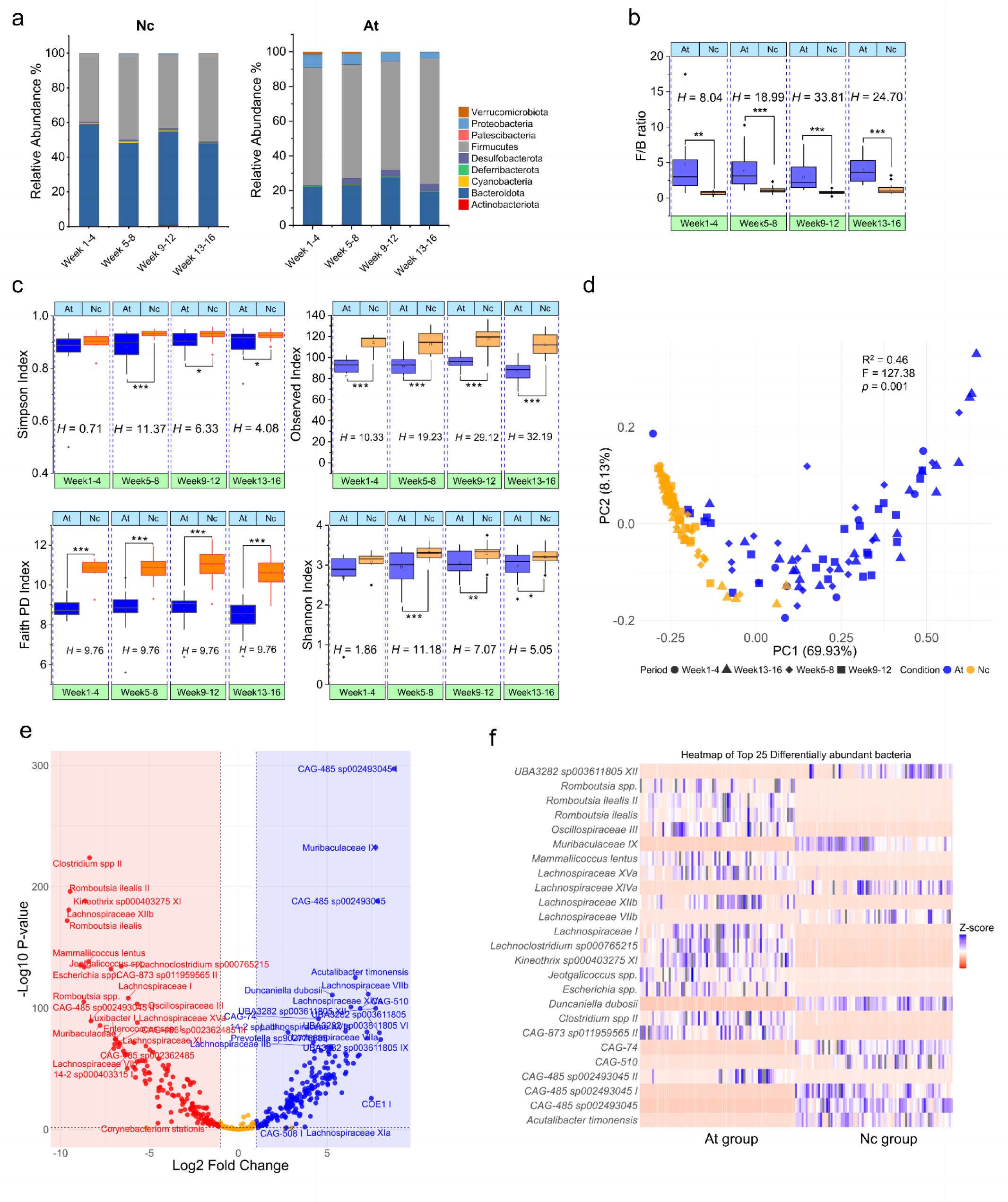
Atherogenic diet causes dysbiosis of gut microbiota. (a) Barchart showing the relative abundance of Nc and At groups in 1-4 weeks, 5-8 weeks, 9-12 weeks and 13-16 weeks. Data are represented in % relative abundance. (b) Boxplot showing the F/B between Nc and At groups. Significance was calculated using Kruskal-Wallis test and value of significance were give as *H* value. \**p* ≤ 0.05; \*\**p* ≤ 0.01; \*\*\**p* < 0.01. (c) Boxplot showing the differences of α-diversity residuals in Nc and At groups using Simpson, Faith, Observed and Shannon indices. (d) Two-dimensional Principal Coordinate analysis using weighted UniFrac distance of Nc and At group base on the gut microbial profile at the species level. PERMANOVA obtained significance with 9999 permutations. (e) Volcano plot of differential abundance analysis determined by DeSeq2 of gut microbiota at the species level. Fold change as a factor of Wald’s test with Benjamini-Hochberg adjusted *p* values is plotted for each species. Significant taxa are represented in different colors (blue, Nc group; red, At group). The orange color represents insignificant tax after the DeSeq2 analysis. (f) Heatmap of the top 25 significant taxa from the DeSeq2 analysis, the gradient was set as low=red, high=blue, mid=white.

The At group (Figure 2a) demonstrated notable change in gut microbiota composition as compare to the gut microbiota of Nc group. The phylum Firmicutes remained the most prevalent during the whole study period. In contrast, the phylum Bacteroidetes exhibited significantly lower relative abundances than the Nc group. Notably, there was an increase (*p* <0.05) in the abundance of the phylum Proteobacteria, which is predominantly associated with pathogenic bacteria^24^, in the At group relative to the Nc group. The Firmicutes/Bacteroidetes (F/B) ratio was significantly elevated (*p* < 0.01) in the At group across all study periods. Specifically, this increase was observed during weeks 1-4 (*H* = 8.04), weeks 5-8 (*H* = 20.98), weeks 9-12 (*H* = 33.81), and weeks 13-16 (*H* = 27.43) (Figure 2b). These findings indicate that the atherogenic diet, which is rich in saturated fatty acids, monounsaturated fatty acids and low in polyunsaturated fatty acids and polysaccharides, modulates the gut microbiota into an enriched Firmicutes and Proteobacteria, but deficient in Bacteroidetes. In contrast, a diet rich in polyunsaturated fatty acids and polysaccharides promotes a balanced gut microbiota, characterized by a lower abundance of Proteobacteria and a more balanced abundance of Bacteroidetes and Firmicutes (Figure S1b).

A significant change in alpha diversity (*p* < 0.01) between Nc group and the At group was observed. A higher alpha diversity in the Nc group was assessed by both the Observed and Faith PD diversity indices (Figure 2c). The disparity in the Observed index is reflected in the number of features obtained from sequencing (Table S2), with the lowest values recorded during weeks 1-4 (*H* = 10.33) and the highest during weeks 13-16 (*H* = 32.19). Similarly, the difference in the Faith PD index was most pronounced during weeks 13-16 (*H* = 9.76). In contrast, there were no significant differences in the Shannon and Simpson indices during weeks 1-4. Nevertheless, relevant differences (*p* < 0.05) were observed from 5-8 -weeks onwards.

To investigate the disparity in gut microbial composition between the Nc and At groups, pairwise PERMANOVA analyses (permutations = 9999) were conducted using the weighted UniFrac distance matrix and a PCoA plot was generated to visualize the clustering patterns based on dietary distinctions. A notable discrepancy in gut microbial composition was identified (R² = 0.46, F = 127.38, p = 0.0001) (Figure 2d). Distinct clusters corresponding to the Nc and At groups were evident. However, a subset of organisms from weeks 9-12 and 13-16 of the At group exhibited clustering with the Nc group. This distinctive clustering of gut microbiota was attributed to variations in dietary composition. Furthermore, distinct phylotypes unique to each group were identified.

Differential abundance analysis at the species level yielded results consistent with the beta diversity analysis. Each group exhibited unique phylotypes, with certain species being abundant in the Nc group but reduced or absent in the At group, and vice versa (Figure 2e). Specifically, *Dunceniella dubosii*, *Acutalibacter timonensis*, CAG-485 sp002493045, CAG-74, and CAG510 were predominantly observed in the Nc group. In contrast, *Romboutsia* spp., *Mammaliicoccus lentus*, *Jeotgalicoccus* spp., *Romboutsia ilealis*, and *Clostridium* spp. were more abundant in the At group (Figure 2f).

### Supplementation of traditional fermented soybean reduces atherogenic diet-induced gut diversity loss and restores a balance gut microbiota akin to normal diet

The supplementation of FSB notably modulated the gut microbiota composition in the atherogenic diet-fed group. Specifically, there was an increase in the relative abundance of the phylum Bacteroidetes and a reduction in the phyla Firmicutes and Proteobacteria in the FSB-supplemented atherogenic diet-fed (AtFSB) group as compared to the At group (Figure 2a). By the end of the 16-week period, the microbial composition of the AtFSB group closely resembled that of the Nc group (Figure 3a). On the contrary, no significant change was observed in the normal chow diet supplemented with FSB (Figure S3a). However, a relative increase in phyla Verricomicrobiota and Firmicutes were observed. The F/B ratio between the Nc group and the AtFSB group showed significant differences (*p* < 0.05) during the first 12 weeks (Figure 3b). However, no significant differences were observed between the two groups during the 13-16 weeks period. A significantly lower F/B ratio ((*p* < 0.01) was observed in AtFSB group as compared to the At group in 5-8 and 13-16 weeks (Figure S2b). Furthermore, no significant differences in alpha diversity, as measured by the Simpson and Shannon indices, were observed between the Nc and AtFSB groups (Figure 3c). Nevertheless, significant differences in Faith PD and the Observed species indices (*p* <0.01) were observed, although the latter showed no significant difference during the last 13-16 weeks period. In the case of At and AtFSB, there was an increase in Shannon, Faith PD, and Observed alpha diversity indices (Figure S2c). These findings suggest that the 13-16 weeks dietary supplementation of FSB in the atherogenic diet regime reduces the abundance of Firmicutes and Proteobacteria and enhances gut microbial diversity similar to the normal diet.

**Figure 3.**
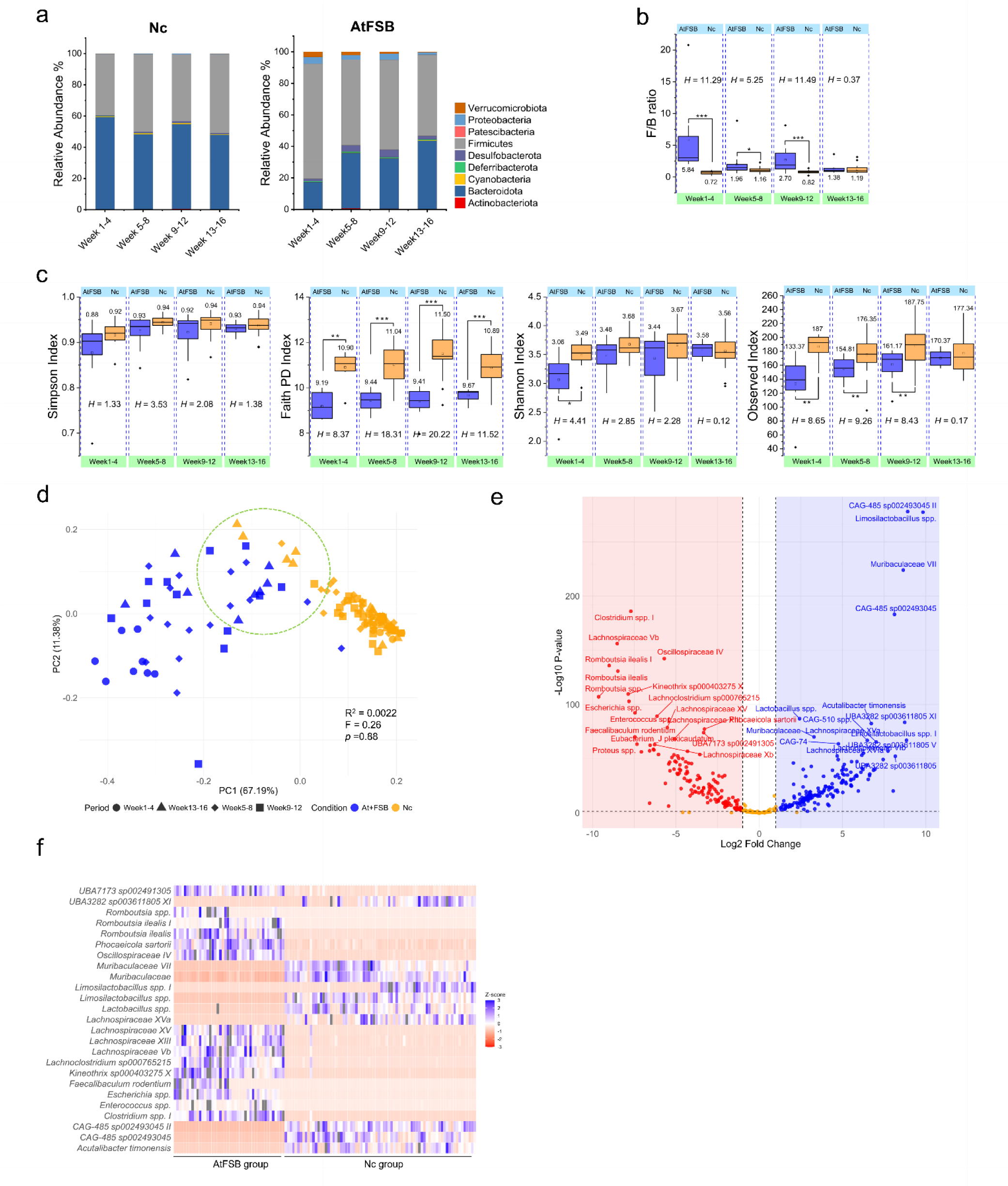
Supplementation of fermented soybean in atherogenic diet group mitigates the loss of diversity and alters the gut microbiota composition, shifting it towards the pattern observed in the normal chow diet group. (a) Barchart showing the relative abundance of Nc and AtFSB groups in 1-4 weeks, 5-8 weeks, 9-12 weeks and 13-16 weeks. Data are represented in % relative abundance. (b) Boxplot showing the F/B between Nc and AtFSB groups. Significance was calculated using Kruskal-Wallis test and the value of significance was give as *H* value. \**p* ≤ 0.05; \*\**p* ≤ 0.01; \*\*\**p* < 0.01. (c) Boxplot showing the differences of α-diversity residuals in Nc and AtFSB groups using Simpson’s, Faith’s, Observed’s and Shannon index. (d) Two-dimensional Principal Coordinate analysis using weighted UniFrac distance of Nc and AtFSB group based on the gut microbial profile at the species level. PERMANOVA obtained significance with 9999 permutations. (e) Volcano plot of differential analysis determined by DeSeq2 of gut microbiota at the species level. Fold change as a factor of Wald’s test with Benjamini-Hochberg adjusted *p* values is plotted for each species. Significant different taxa are represented in different colors (blue, Nc group; red, AtFSB group). The orange color represents insignificant tax after the DeSeq2 analysis. (f) Heatmap of the top 25 significant taxa from the DeSeq2 analysis, the gradient was set as low=red, high=blue, mid=white.

Pairwise PERMANOVA analysis (9999 permutations) utilizing the weighted UniFrac distance matrix between the Nc and AtFSB groups indicated no significant differences in microbial composition (R² = 0.002, F = 0.26, *p* = 0.88) as indicated by the clusering patterns in Figure 3d. Notably, microbial communities from weeks 13-16 of the Nc group cluster with those from weeks 5-8, 9-12, and 13-16 of the AtFSB group. Consistent findings were observed between At and AtFSB (R² = 0.011, F = 1.42, *p* = 0.21) (Figure S2d) and between Nc and NcFSB (R² = 0.008, F = 1.06, *p* = 0.35) (Figure S3d).

Differential abundance analysis revealed distinct microbial compositions between the Nc and AtFSB groups (Figure 3e). Specifically, the family Muribaculaceae, CAG-485 spp., *Limosilactobacillus* spp, *Lactobacillus* spp., and *Acutalibacillus timonensis* were more abundant in the Nc group, whereas *Clostridium* spp. I, *Romboutsia* spp., *Phocaeicola sartori*, *Faecalibaculum rodentium*, *Escherichia* spp., and the families Lachnospiraceae and Oscillospiraceae were more prevalent in the AtFSB group (Figure 3f). *Bacillus* spp., *Coprocola* sp9000364165, *Lawsonibacter* sp00492175, UBA3282 sp00311805 were differentially abundant in AtFSB than the At (Figures S2e and 2f). In addition, *Angelakisella massiliensis*, *Eubacterium* sp000687695, *Acetalifactor* sp011959105, *Merdisoma* sp011959465, *Odoribacter laneus*, and the family Lachnospiraceace were dominant in NcFSB than the Nc group (Figure S3f).

### Diet-induced alteration in the core microbiota of the gut

The gut core microbiota was analyzed using QIIME2, the core microbiome may be defined as taxa observed in at least half of the samples. The composition was assessed across fractions of samples from 0.5 to 1, at intervals of 0.05 (totaling 11 intervals). Among the 299 core taxa identified across four groups (Figure 4a), 63 taxa were common to all groups (Figure 4b), while 30 taxa were unique to the Nc group (Figure 4c), 12 to the Atherogenic group (Figure 4d), 41 to the NcFSB group (Figure 4e), and 31 to the AtFSB group (Figure 4f). In the Nc group, the predominant core taxa included CAG-485 sp002493045 (11.16%), *Clostridium disporicum* (4.88%), *Turicibacter* spp. (4.41%), *Muribaculum gordoncarteri* II (3.15%), *Faecalibaculum rodentium* (2.64%), *Ligilactobacillus* spp. (2.26%), *Muribaculum gordoncarteri* I (1.77%), *Duncaniella dubosii* (1.52%), *Phocaeicola sartorii* (1.42%), and *Desulfovibrio trichonymphae* (1.32%). In the NcFSB group, the dominant core taxa were CAG-485 sp002493045 (12.41%), *Turicibacter* spp. (5.24%), *Ligilactobacillus* spp. (4.30%), *Muribaculum gordoncarteri* II (2.54%), *Duncaniella dubosii* (2.52%), *Desulfovibrio trichonymphae* (2.36%), Muribaculaceae II (2.34%), *Faecalibaculum rodentium* (2.07%), *Muribaculum gordoncarteri* I (1.82%), and *Clostridium disporicum* (1.73%).

**Figure 4.**
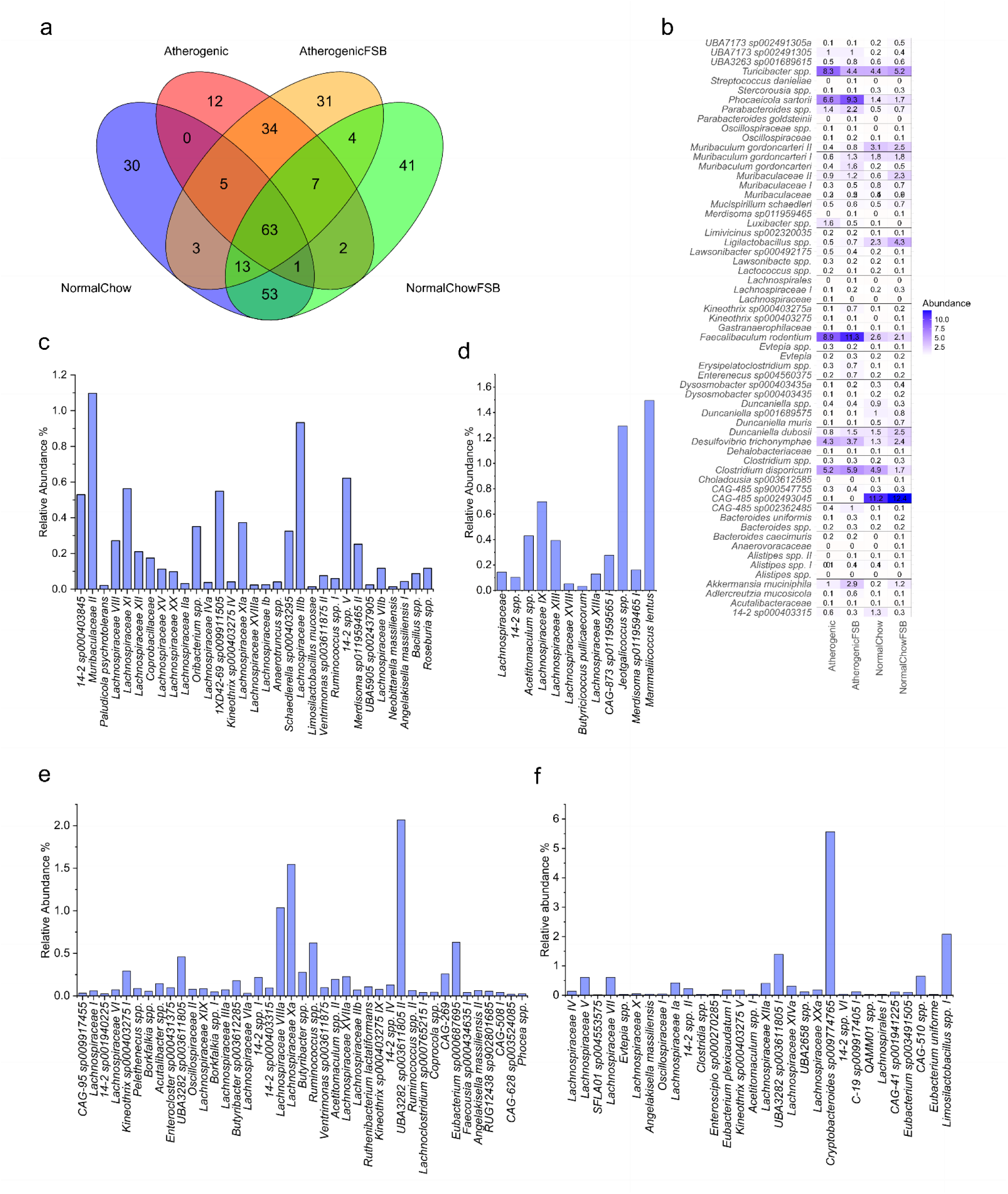
Core microbiome of gut mice on different dietary conditions. (a) Venn diagram showing the core microbiome of Nc, At, NcFSB and AtFSB group. (b) Heat map of 63 common core bacteria present in Nc, At, NcFSB and AtFSB groups. The value within the cell represents the % relative abundance of each taxon. (c) Bar Chart of the core bacteria present only in the Nc group. (d) Bar Chart of the core bacteria present only in the At group. (e) Bar Chart of the core bacteria present only in the NcFSB group. (f) Bar Chart of the core bacteria present only in the AtFSB group.

In the At group, the most prevalent core taxa included *Faecalibaculum rodentium* (8.88%), *Turicibacter* spp. (8.33%), *Phocaeicola sartorii* (6.56%), *Clostridium disporicum* (5.24%), *Desulfovibrio trichonymphae* (4.26%), *Luxibacter* spp. (1.57%), *Parabacteroides* spp. (1.35%), *Akkermansia muciniphila* (1.03%), UBA7173 sp002491305 (1.00%), and Muribaculaceae II (0.91%). In the AtFSB group, the dominant core taxa were *Faecalibaculum rodentium* (11.31%), *Phocaeicola sartorii* (9.28%), *Clostridium disporicum* (5.93%), *Turicibacter* spp. (4.40%), *Desulfovibrio trichonymphae* (3.71%), *Akkermansia muciniphila* (2.86%), *Parabacteroides* spp. (2.19%), *Muribaculum gordoncarteri* (1.59%), *Duncaniella dubosii* (1.49%), and *Muribaculum gordoncarteri* I (1.32%).

The divergence in core taxa between the Nc and At group was markedly pronounced. Fermented soybean supplementation in the normal diet (NcFSB group) increased the abundance of the *Muribaculum* genus and the Muribaculaceae family, accompanied by a reduction in the prevalence of *Clostridium disporicum*. In the AtFSB group, there was a notable augmentation in the abundance of *Muribaculum* genus, *Phocaeicola sartorii*, *Faecalibaculum rodentium*, and *Akkermansia muciniphila*, coupled with a decrease in *Turicibacter* spp. (Figure 4b). These findings suggest that the elevated presence of the *Muribaculum* genus and Muribaculaceae family within the core taxa may play a crucial role in sustaining gut microbiota diversity and overall gut health.

### Functional prediction with PICRUSt2 revealed changes in metabolic activities

PICRUSt2 analysis identified 4,843 KEGG Orthologs (KOs) differentiating Nc and At groups, 4,963 KOs between Nc and AtFSB, 3,708 KOs between Nc and NcFSB, and 5,056 KOs between At and AtFSB. All major functional pathways were more enriched in the At group compared to the Nc group (Figure 5a). Upon FSB supplementation in the atherogenic diet, a notable reduction was observed in functional categories such as amino acid metabolism, carbohydrate metabolism, lipid metabolism, energy metabolism, and the metabolism of cofactors and vitamins, as well as in unclassified metabolic processes (Figure 5d). However, the metabolism of terpenoids and polyketides increased in AtFSB. The decline in amino acid and carbohydrate metabolisms in the AtFSB group closely resembled the levels seen in the Nc group (Figure 5b). Supplementation with FSB in the normal diet led to a decrease in metabolisms, with marked reduction in the nucleotide metabolism, and metabolism of terpenoid and polyketide (Figure 5c).

**Figure 5.**
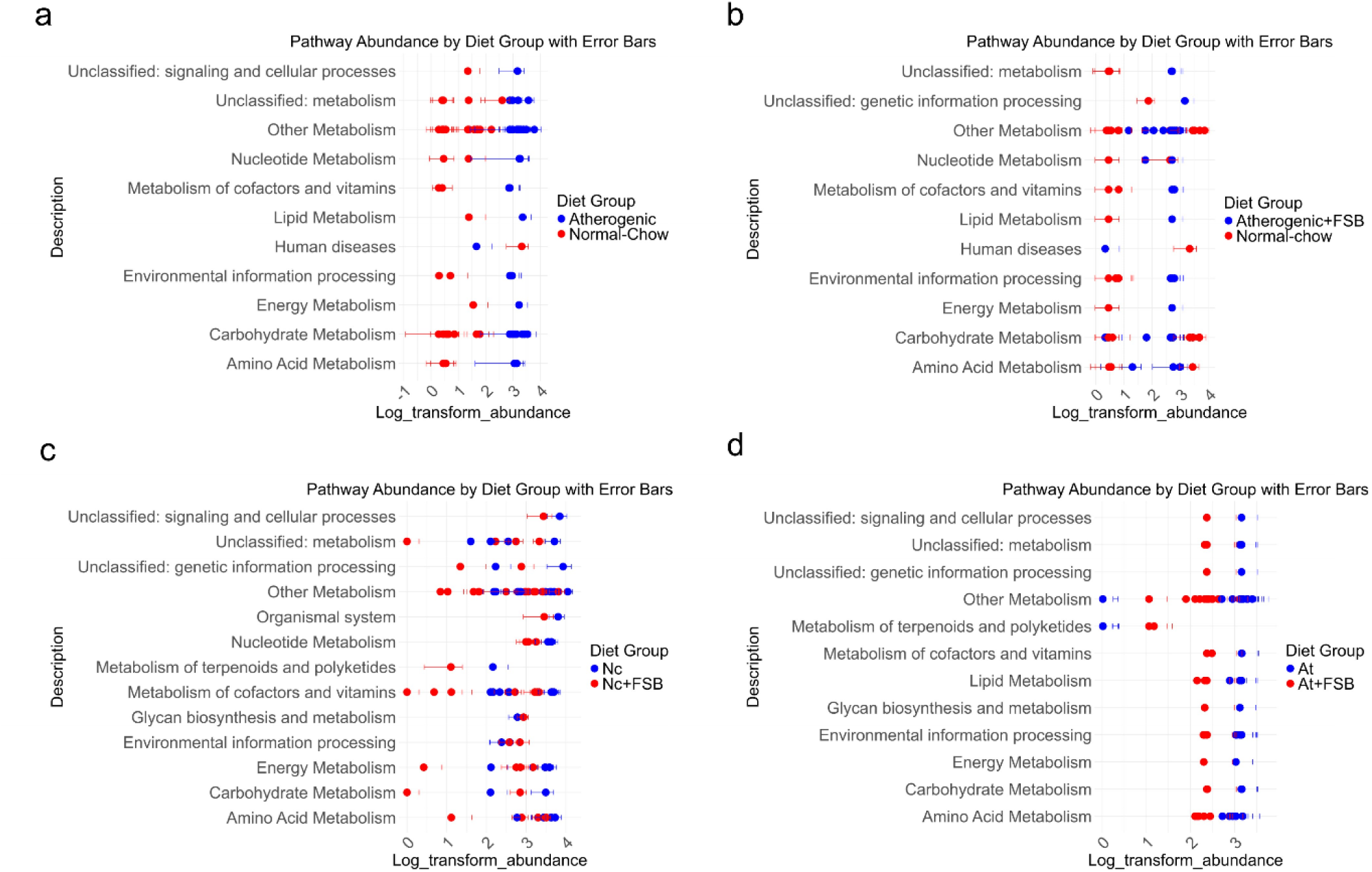
Functional prediction of gut microbiota in PICRUSt 2 using 16S-V4 hypervariable region as a marker. (a) A functional categories abundance graph between Nc and At with error bar. (b) A functional categories abundance graph between Nc and AtFSB with error bar. (c) A functional categories abundance graph between Nc and NcFSB with error bar. (d) A functional categories abundance graph between At and AtFSB with an error bar.

## Discussion

The atherogenic diet in this study has 20% crude fat, out of which more than 87 % is composed of saturated fatty acid, roughly 6% of monounsaturated fatty acid, and the rest 7 % with polyunsaturated fatty acid. Of the total carbohydrates present in the diet, 82.7% is comprised of simple sugar and starch contributes only 17.3% in the atherogenic diet. Whereas, the normal chow diet has 4.1% crude fat, 60.54% polyunsaturated fatty acid, 16.82% saturated fatty acid, and 22.64% monounsaturated fatty acid. In the normal chow diet composition, 89.12% comprise of starch and 10.88% with simple sugars of the total carbohydrates content. The disparity in the diet composition causes a significant change in the gut microbiota composition between the Nc group and the At group. Increased abundance of Firmicutes and reduced Bacteroidetes was reported in high-fat diet-fed mice^25,26^ and also increased F/B ratio^27^ and reduced alpha diversity^28^ leading to gut dysbiosis, which were observed in the At group. Increase in phylum Proteobacteria associated inflammatory and dysbiosis^29^ and phylum Verrucomicrobia^30^ is linked with beneficial gut health such as gut barrier function and inflammation control was reduced in At group. Thus, a diet rich in saturated fatty acid, monounsaturated fatty acid and less polyunsaturated fatty acid combined with higher simple sugars and lower starch causes dysbiosis in gut microbiota, which can lead to various health issues.

Elevated levels of total cholesterol and low-density lipoprotein (LDL) are the major risk factors for cardiovascular diseases. Increased levels of LDL-C posed a risk of myocardial infarction and atherosclerosis^31,32^. Management of LDL-C plays a crucial role in lowering the CVD risk^33^. In our study, the At group has a higher total cholesterol and LDL-C (Figures 1d and 1e), coupled with a higher gene expression level of pro-inflammatory markers IL-1β^34^, CD4^35^, ICAM-1^36^. In addition, the presence of atherosclerotic lesions in the intima layers of the arch of the aorta (Figures 1a and 1b) confirmed the atherosclerosis in the At group. Since the gut is permeable (Figures 1g and 1h) because of the week tight junction (Figure 1i), lipopolysaccharides (LPS) from the gram-negative bacteria might have aggravated the atherosclerotic condition^37^. In the functional prediction (Figure S4a), the prostaglandin reductase-3 pathway involved in the arachidonic acid metabolism is more abundant in the At group, which, coupled with higher serum endotoxin level due to the dysfunctional tight junction and increased level of pro-inflammatory cytokines^38^, might have resulted in the development of atherosclerosis^39^.

Supplementation of FSB ameliorated the atherogenic diet-induced diversity lost by increasing the abundance of Bacteroidetes and reducing the abundance of Firmicutes and Proteobacteria (Figure S2a). Lower F/B ratios were also observed in FSB supplemented diet group (Figure S2b; Figure S3b). Alpha diversity increased significantly when supplemented with FSB in the AtFSB group (Figure 3c, Figure S2c), however, the β-diversity seen between Nc and AtFSB was insignificant (*p*=0.88) (Figure 3d). Alteration in the core microbiome was also observed in FSB supplemented groups (Figure 4). A study on Thau Nau, a traditional fermented soybean of Thailand, revealed that it is a potential source of SCFAs^40^. For instance, SCFAs-producing bacteria such as *Angelakisella massiliensis*^41^, *Eubacterium* spp.^42^, *Limosilactobacillus* spp.^43^, etc., were abundant in the groups supplemented with FSB. All these may influence the reshaping of the dysbiosis gut microbiota to a near-eubiotic state.

Additionally, FSB reduced the atherosclerotic lesion by lowering the gene expressions of pro-inflammatory markers such as IL-1β, CD4, and ICAM-1 (Figures 1a, 1b and 1c). No significant change was found in the total cholesterol and LDL-C serum in the FSB-supplemented group. FSB improves gut permeability by increasing the expression of tight junction genes (Figure 1i), thereby decreasing the gut permeability and reducing the endotoxin level in the serum (Figures 1g and 1h). The effect of FSB can also be further seen in the functional prediction (Figures S4c and S4d). The metabolism of terpenoids and polyketides was abundant only in the group supplemented with FSB. Terpenoids and polyketides are known for their antimicrobial activity and cytotoxic activities^44^ and crucial for host metabolism and defence mechanisms^45^. The pathway for the enzymes farnesyl diphosphate synthase, plipastatin/fengycin lipopeptides synthetase A/C were significantly abundant in FSB-supplemented group. These pathways are involved in the production of bioactive metabolites isoprenoids^46^, and lipopeptides such as plipastatin^47^, fengycins^48^, which are known bioactive compounds that exert antimicrobial, antifungal and various other health benefits that improve gut function, and maintain the gut microbial balance. Reduce abundance in pathogenic phylum Proteobacteria and returning of near-eubiosis state may be due to the abundance of bioactive metabolites pathways. In our study, we did all the animal experiments using the wild-type mice C57BL6/N Tac only, and we tried to induce atherosclerosis by feeding an atherogenic diet only. Using standard knock-out models such as ApoE-/-and LDL-/- for atherosclerosis may provide mechanistic insights behind fermented soybean modulation of gut microbiota, gut health and atherogenesis.

## Conclusion

A diet rich in saturated and monounsaturated fatty acids, monosaccharides, and disaccharides, cholesterol and low in polyunsaturated fatty acids and polysaccharides causes loss of gut microbial diversity, dysbiosis, and gut barrier integrity, and poses a higher risk of developing atherogenic lesions. Long-term consumption of fermented soybeans rich in bioactive compounds reverses gut microbiota alterations, improves gut health and reduces lesion formation. Our study underscores the significance of traditional fermented soybean in maintaining gut microbiota diversity and restoring gut health against dietary-induced dysbiosis, offering important insights into its role in modulating atherogenesis in a gut microbiota-dependent manner.

## Data availability

The raw sequence reads data from 16S metagenomic sequencing that were generated in and support the findings of this study have been deposited and are publicly available in the NCBI Sequence Read Archive at https://www.ncbi.nlm.nih.gov/bioproject/?term=(PRJNA1143515)), accession reference PRJNA1143515.

## Acknowledgments

The authors thank the Publication & Intellectual Property Rights Committee, CSIR-NEIST, Jorhat for approving the manuscript for publication (Manuscript No: CSIR-NEIST/PUB/2024/075, dated 14-10-2024). We thank the Centre for Pre-clinical Study (CPS), CSIR-NEIST, for supporting the research by providing animal husbandry facilities to conduct animal-related experiments. Special acknowledgment is extended to Mr. Prashanta Bora, Laboratory Attendant, CPS, CSIR-NEIST, for his invaluable assistance during the animal experimentation phases. We would like to thank the Director, CSIR-NEIST, for providing the overall support and facility to carry out the work.

## CRediT Roles

**R.W.:** Conceptualization, Data curation, Formal analysis, Funding acquisition, Investigation, Methodology, Project administration, Software, Resources, Supervision, Writing – original draft, Writing – review & editing. **H.K.B.:** Methodology, Resources, Writing – review & editing. **M.G.S.:** Data curation, Formal analysis, Funding acquisition, Investigation, Methodology, Software, Visualization, Writing – original draft, Writing – review & editing.

## Supplementary figures

**Figure S1.**
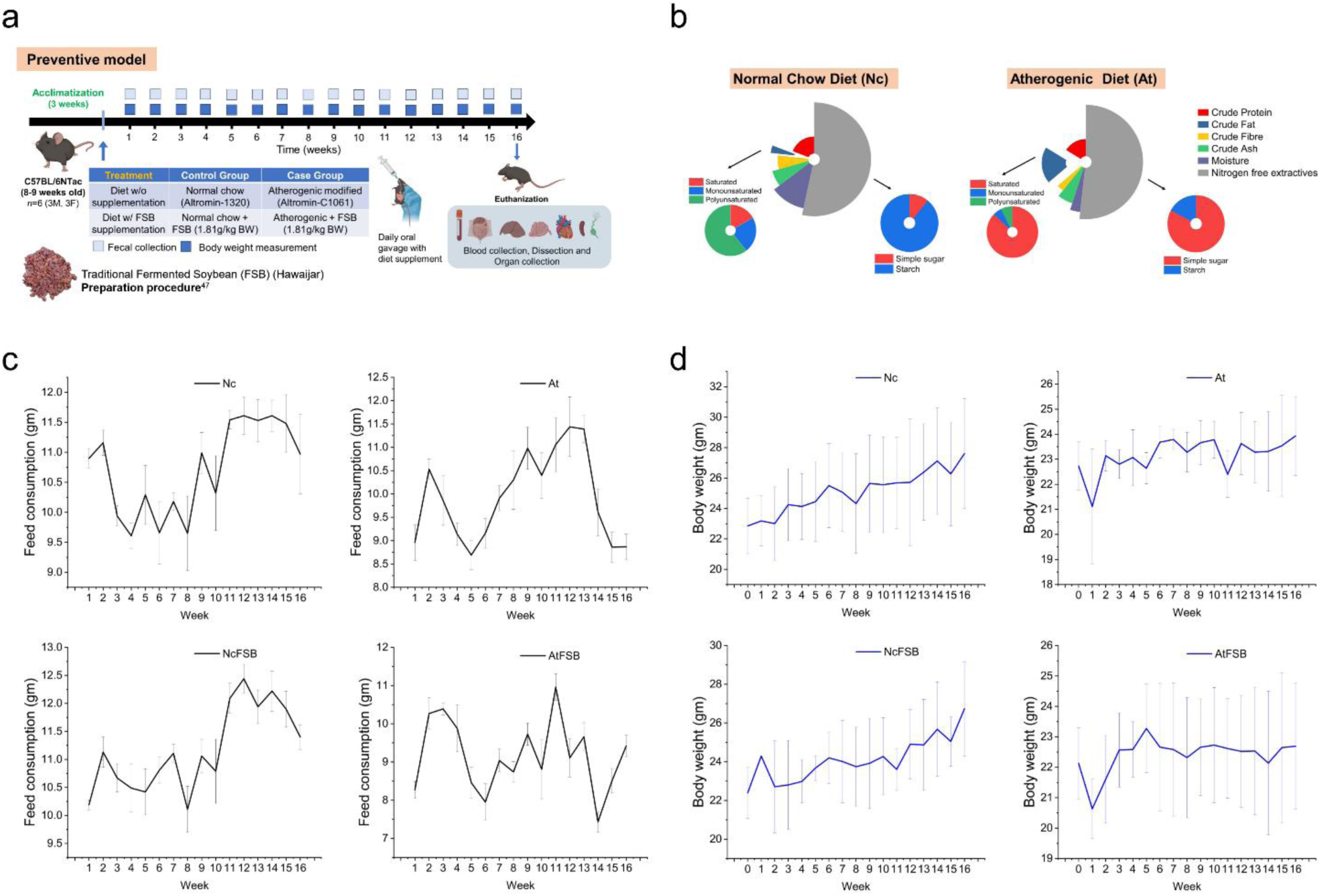
Change in feed consumption and body weight during the 1-16 week period. (a) Schematic diagram of experimental design fed with traditional fermented soybean (Hawaijar). (b) Diet composition of standard normal chow diet and atherogenic diet. (c) Feed consumption pattern of different experimental groups. (d) Change in body weight of animals of different diet regimes from 0 to 16 weeks.

**Figure S2.**
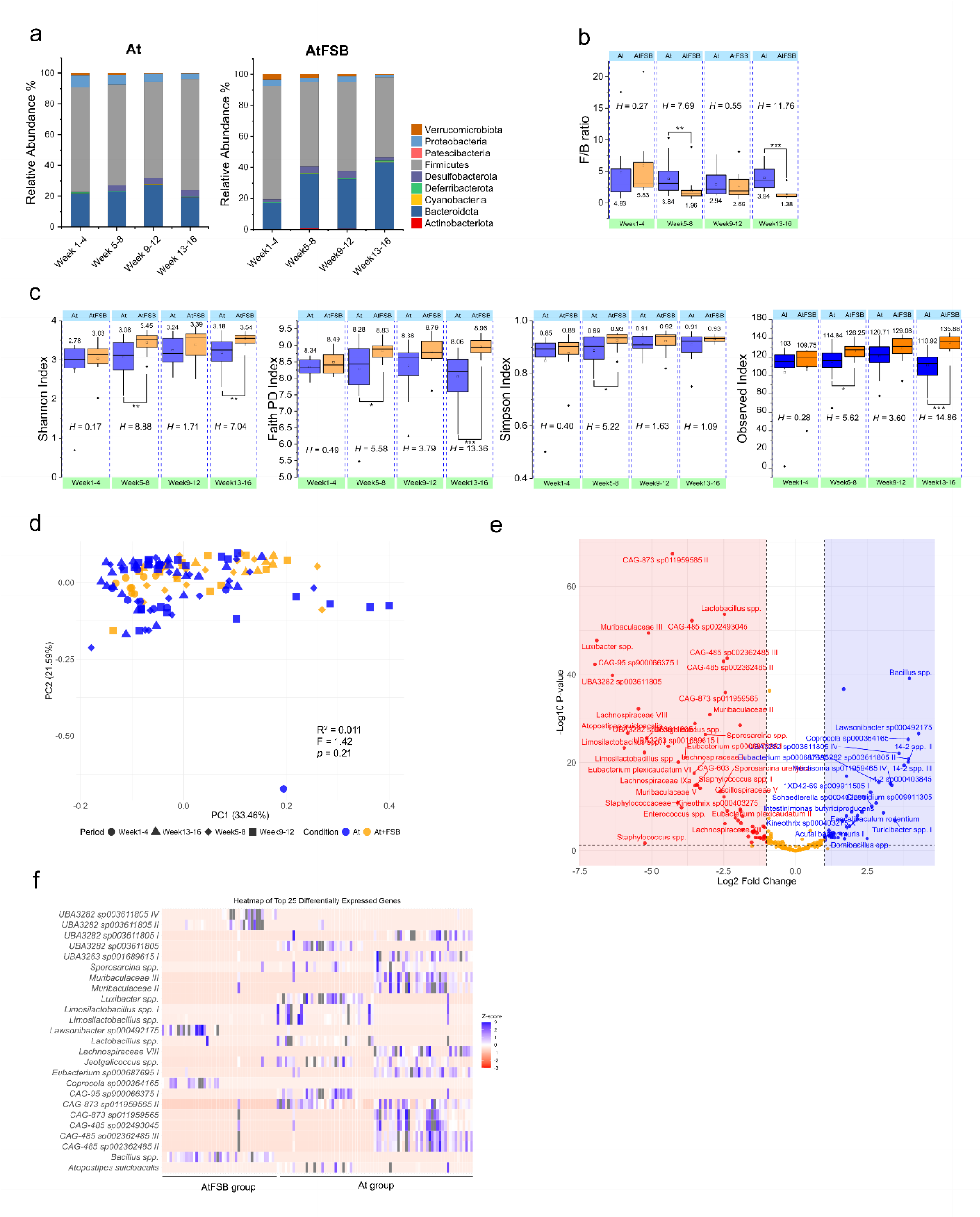
Difference in the gut microbiota composition between At group and AtFSB group. (a) Barchart showing the relative abundance of At and AtFSB groups in 1-4 weeks, 5-8 weeks, 9-12 weeks and 13-16 weeks. Data are represented in % relative abundance. (b) Boxplot showing the F/B between At and AtFSB groups. Significance was calculated using the Kruskal-Wallis test, and the strength of significance were given as H values. (c) Boxplot showing the differences of α-diversity residuals in At and AtFSB groups using Simpson, Faith, Observed, and Shannon indices. (d) Two-dimensional Principal Coordinate analysis using weighted UniFrac distance of At and AtFSB group based on the gut microbial profile at the species level. PERMANOVA obtained significance with 9999 permutations. (e) Volcano plot of differential analysis of gut microbiota at the species level determined by DeSeq2. Fold change as a factor of Wald’s test with Benjamini-Hochberg adjusted *p* values was plotted for each species. Significant different taxa are represented in different colors (blue, AtFBS group; red, At group). The orange color represents insignificant tax after the DeSeq2 analysis. (f) Heatmap of the top 25 significant taxa from the DeSeq2 analysis, the gradient was set as low=red, high=blue, mid=white.

**Figure S3.**
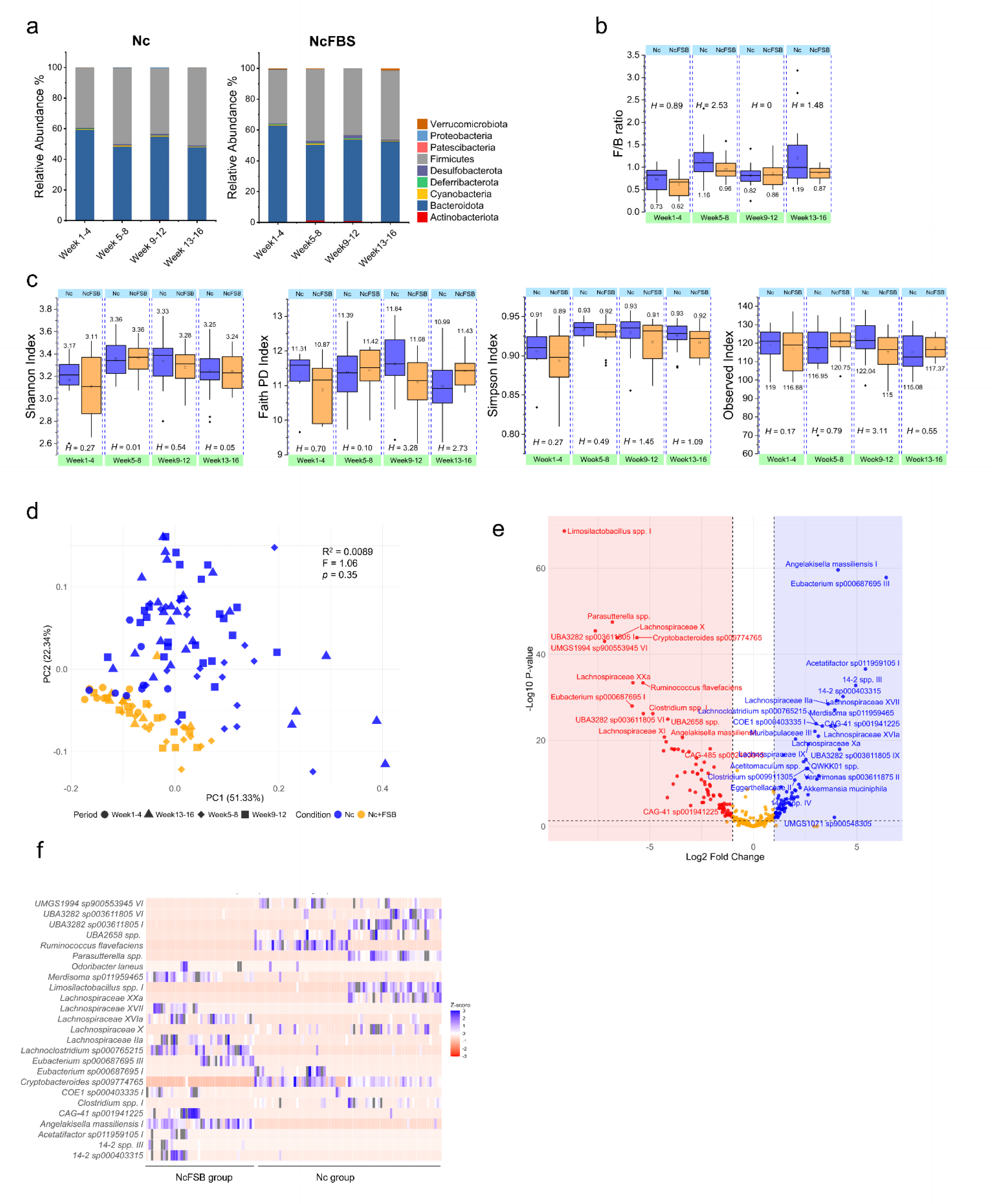
Difference in the gut microbiota composition between Nc group and NcFSB group. (a) Barchart showing the relative abundance of Nc and NcFSB groups in 1-4 weeks, 5-8 weeks, 9-12 weeks and 13-16 weeks. Data are represented in % relative abundance. (b) Boxplot showing the F/B between Nc and NcFSB groups. Significance was calculated using Kruskal-Wallis test and values of significance were give as *H* values. (c) Boxplot showing the differences of α-diversity residuals in Nc and NcFSB groups using Simpson, Faith, Observed and Shannon indices. (d) Two-dimensional Principal Coordinate analysis using weighted UniFrac distance of At and AtFSB group based on the gut microbial profile at the species level. PERMANOVA obtained significance with 9999 permutations. (e) Volcano plot of differential analysis of gut microbiota at the species level determined by DeSeq2. Fold change as a factor of Wald’s test with Benjamini-Hochberg adjusted *p* values was plotted for each species. Significant different taxa are represented in different colors (blue, NcFBS group; red, Nc group). The orange color represents insignificant tax after the DeSeq2 analysis. (f) Heatmap of the top 25 significant taxa from the DeSeq2 analysis, the gradient was set as low=red,high=blue, mid=white.

**Figure S4.**
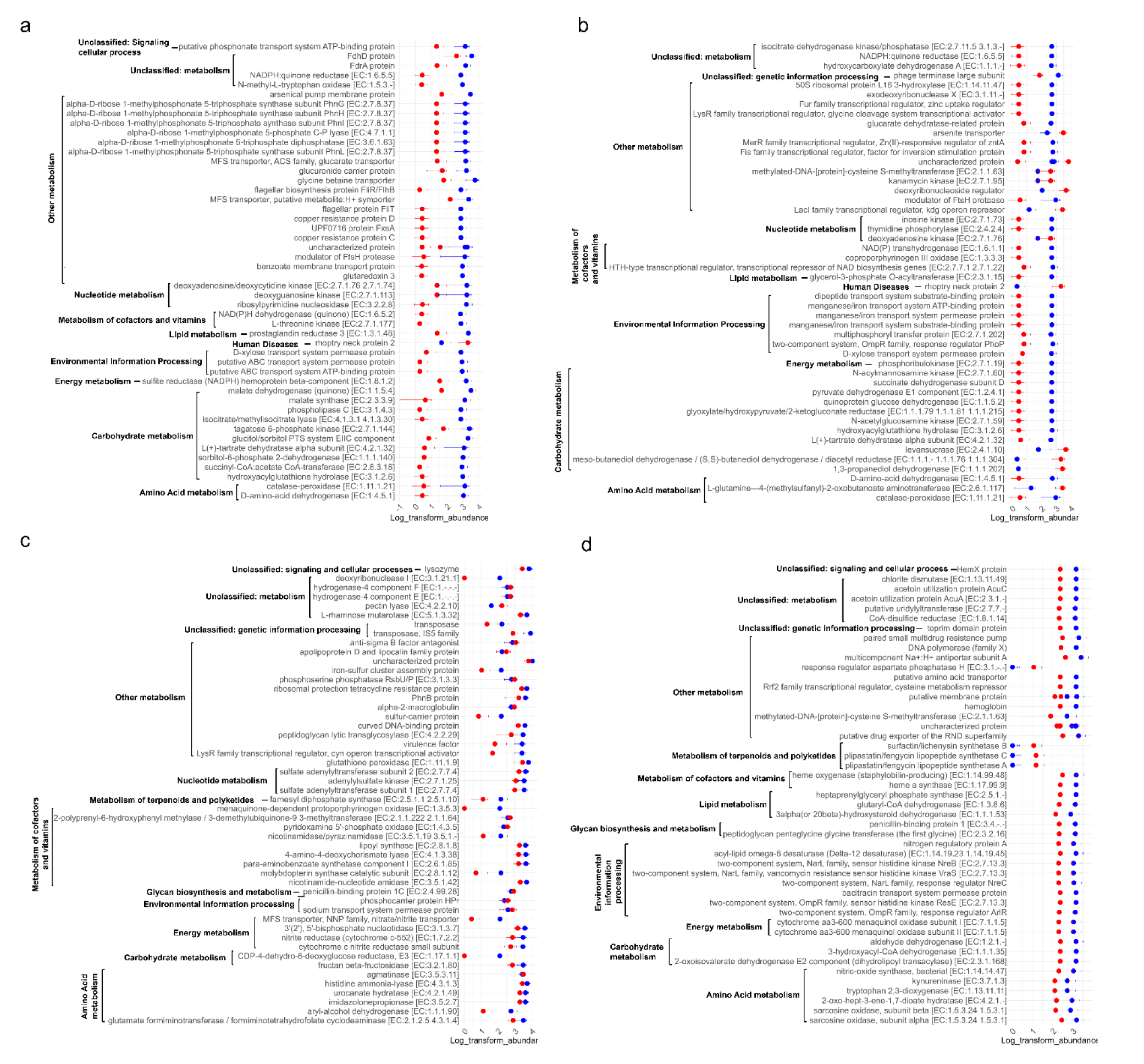
Functional prediction of gut microbiota using 16S-V4 hypervariable region as a marker in PICRUSt 2. (a) A pathway abundance graph between Nc (red) and At (blue) with error bar. (b) A pathway abundance graph between Nc (red) and AtFSB (blue) with error bar. (c) A pathway abundance graph between Nc(blue) and NcFSB(red) with error bar. (d) A pathway abundance graph between At (blue)and AtFSB(red) with an error bar.

**Figure S5.**
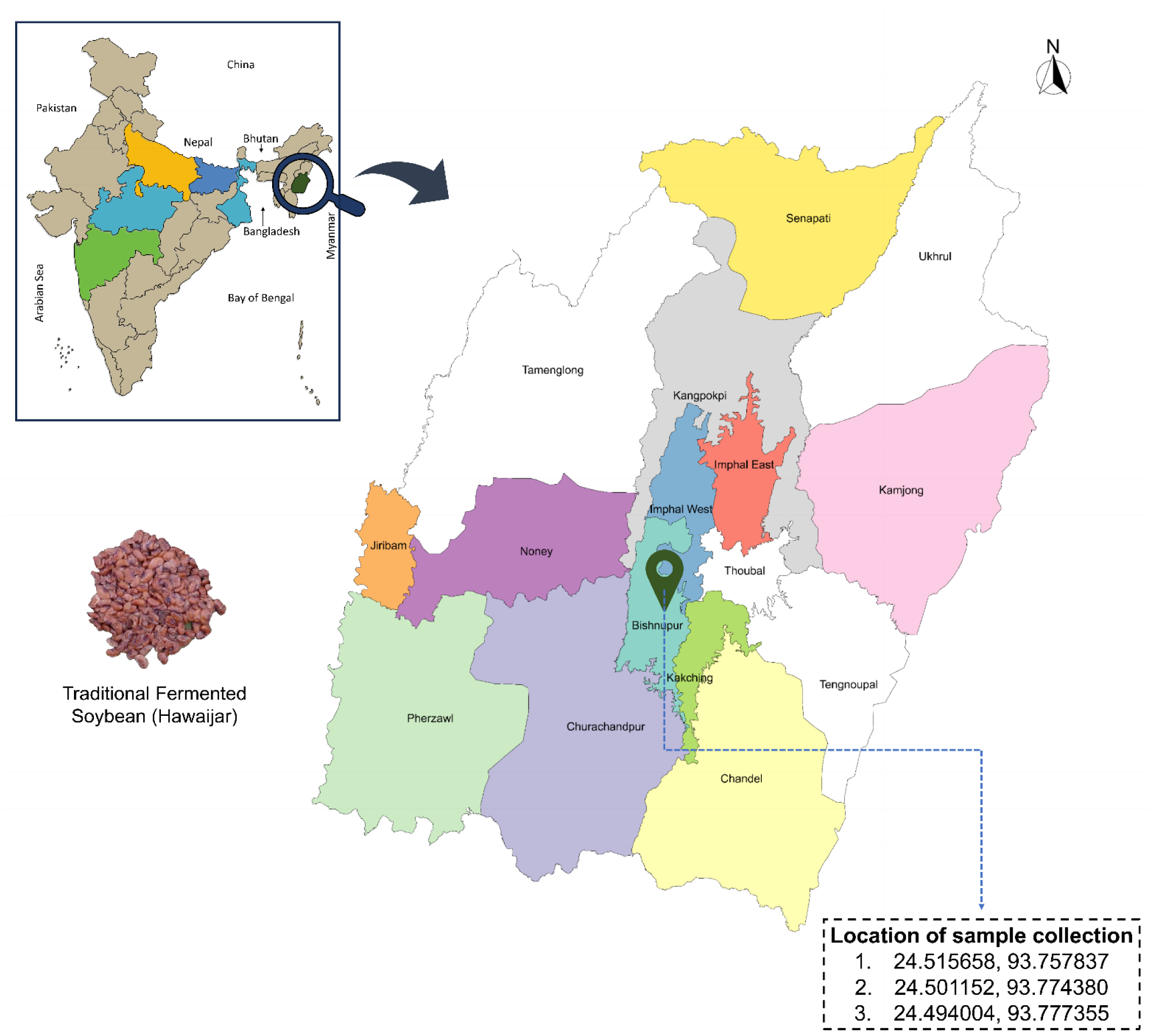
Sample collection sites for traditional fermented soybean (Hawaijar) in Bishnupur district, Manipur.

## Supplementary Tables

**Table S1.**
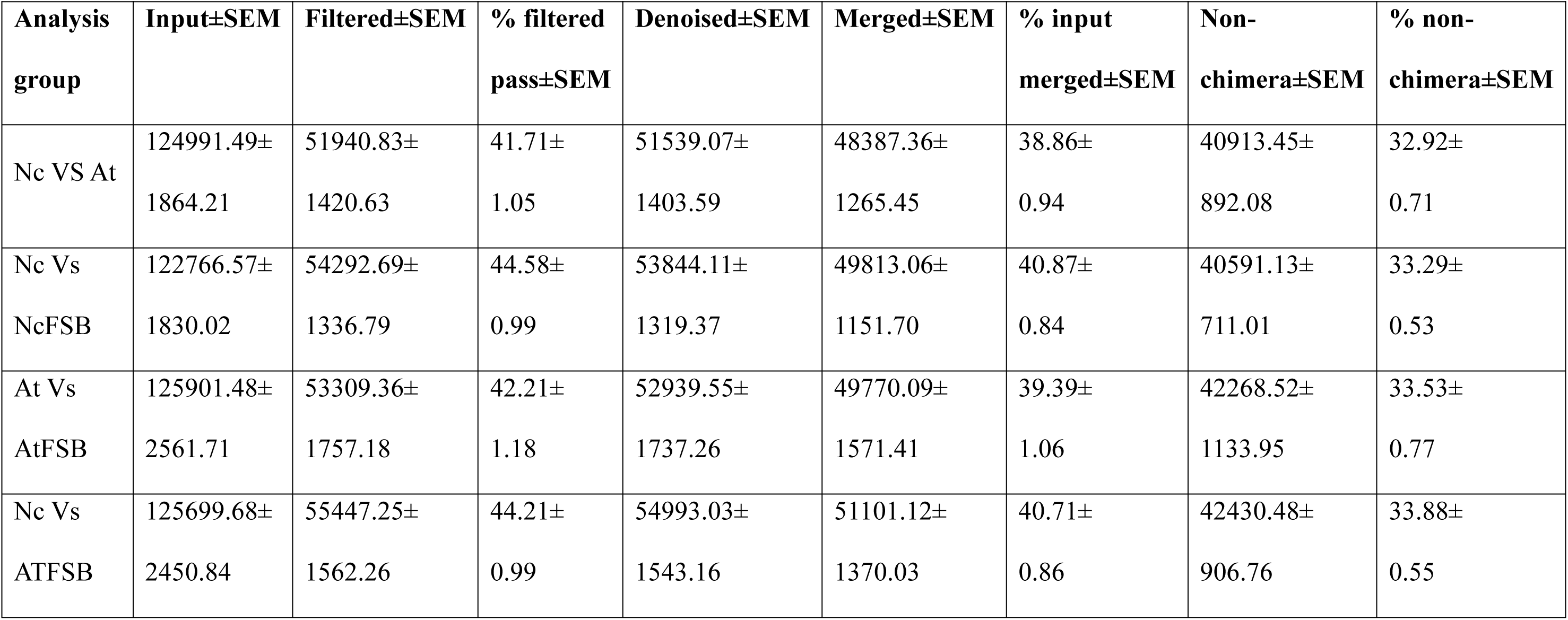
Summary statistics of reads after filtering and denoising using DADA2 plugins.

**Table S2.**
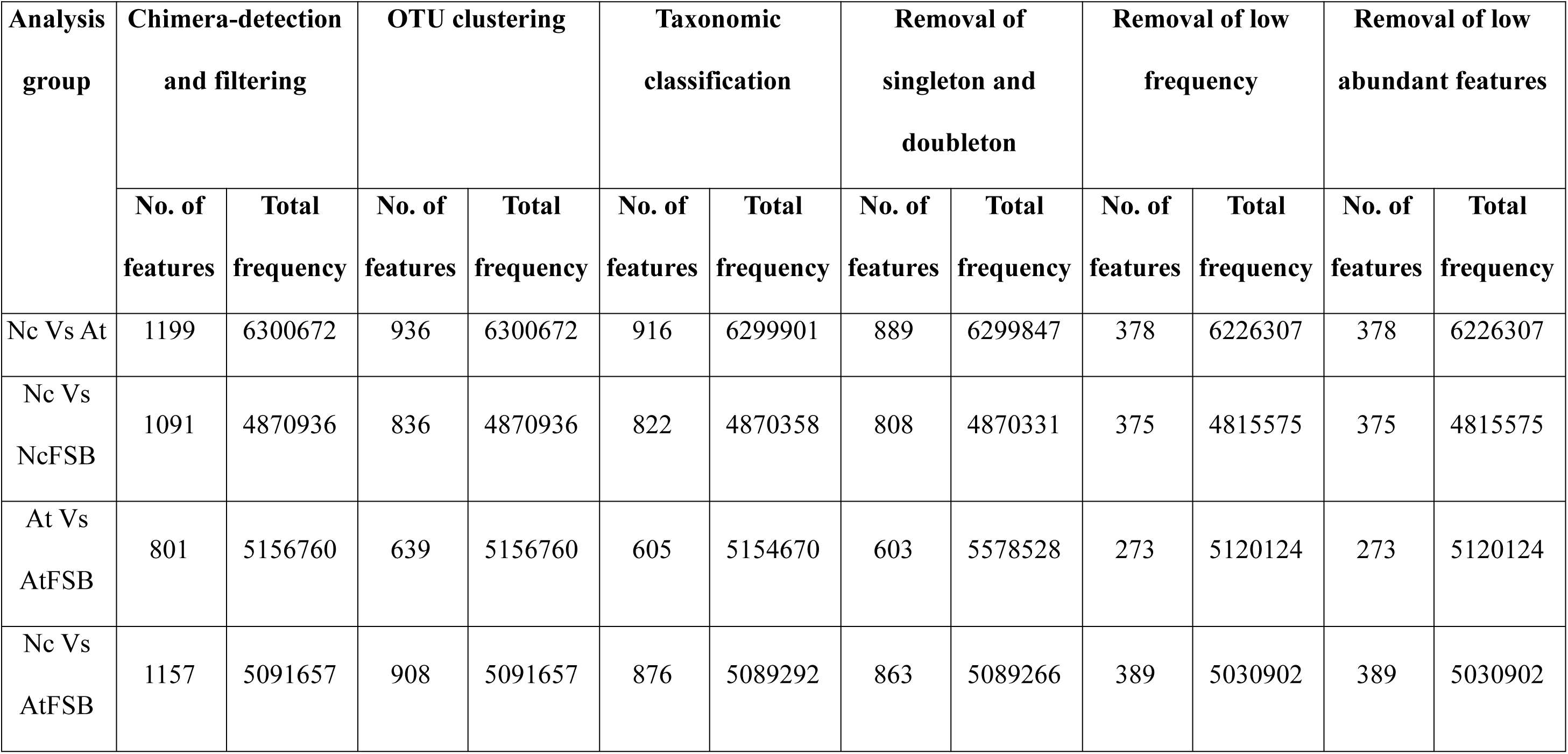
Filtering of feature and frequency during QIIME2 analysis.

| Analysis group | Chimera-detection and filtering |  | OTU clustering |  | Taxonomic classification |  | Removal of singleton and doubleton |  | Removal of low frequency |  | Removal of low abundant features |  |
| --- | --- | --- | --- | --- | --- | --- | --- | --- | --- | --- | --- | --- |
|  | No. of features | Total frequency | No. of features | Total frequency | No. of features | Total frequency | No. of features | Total frequency | No. of features | Total frequency | No. of features | Total frequency |
| Nc Vs At | 1199 | 6300672 | 936 | 6300672 | 916 | 6299901 | 889 | 6299847 | 378 | 6226307 | 378 | 6226307 |
| Nc Vs<br>NcFSB | 1091 | 4870936 | 836 | 4870936 | 822 | 4870358 | 808 | 4870331 | 375 | 4815575 | 375 | 4815575 |
| At Vs<br>AtFSB | 801 | 5156760 | 639 | 5156760 | 605 | 5154670 | 603 | 5578528 | 273 | 5120124 | 273 | 5120124 |
| Nc Vs<br>AtFSB | 1157 | 5091657 | 908 | 5091657 | 876 | 5089292 | 863 | 5089266 | 389 | 5030902 | 389 | 5030902 |

**Table S3.**
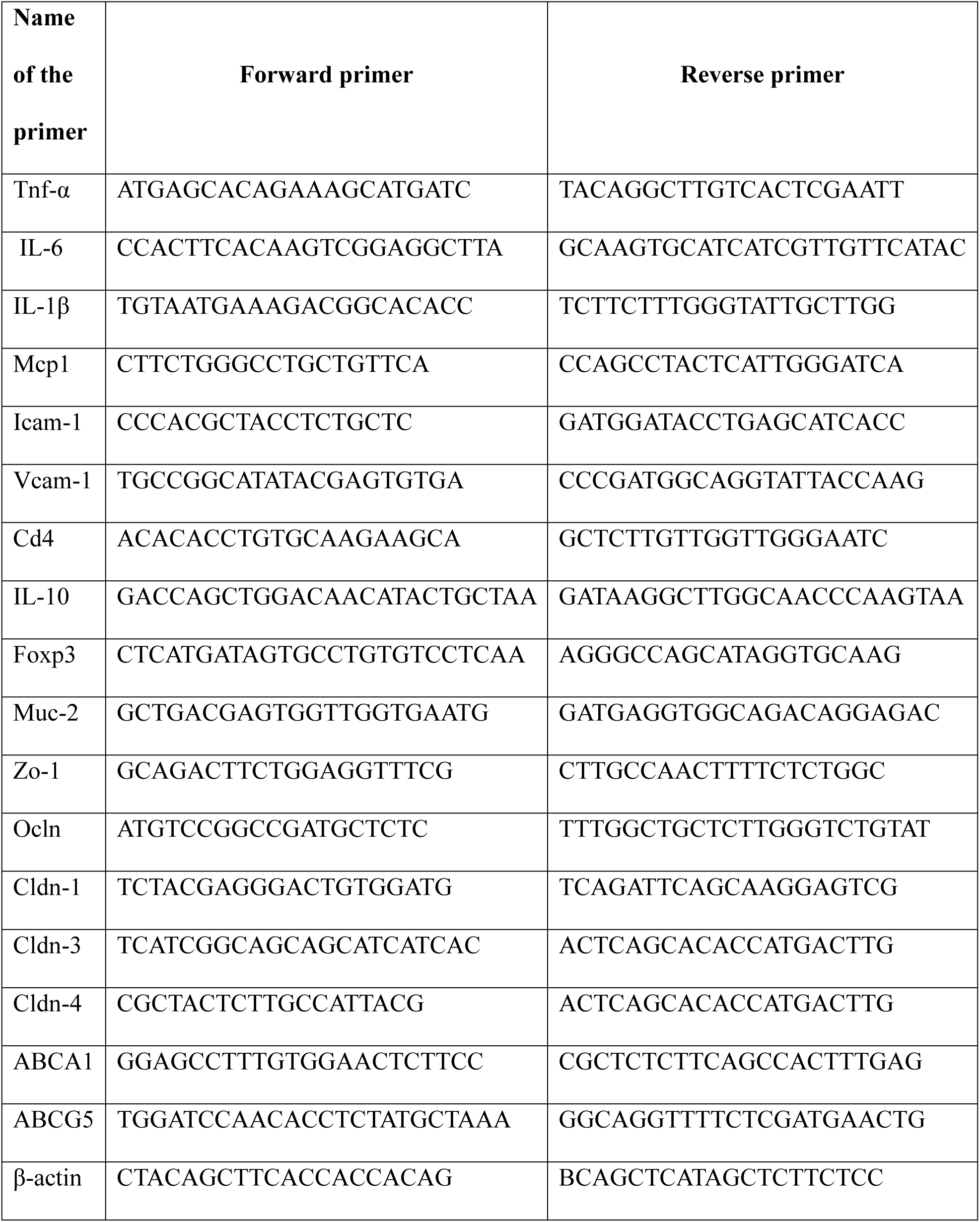
List of qPCR primer name and sequence.

